# Molecular aspects of multivalent engagement between Syk and FcεRIγ

**DOI:** 10.1101/469148

**Authors:** Timothy Travers, William Kanagy, Elton Jhamba, Byron Goldstein, Diane S. Lidke, Bridget S. Wilson, S. Gnanakaran

**Author notes:** Current address: New Mexico Consortium and Pebble Labs Inc., Los Alamos, NM 87544.

## Abstract

Syk/Zap70 family kinases are essential for signaling via multichain immune-recognition receptors such as the tetrameric (αβγ2) FcεRI The simplest model assumes that Syk activation occurs through *cis* binding of its tandem SH2 domains to dual phosphotyrosines within immunoreceptor tyrosine-based activation motifs of individual γ chains. In this model, Syk activity is modulated by phosphorylation occurring between adjacent Syk molecules docked on γ homodimers and by Lyn molecules bound to FcεRIβ. However, the mechanistic details of Syk docking on γ homodimers are not fully resolved, particularly the possibility of *trans* binding orientations and the impact of Y130 autophosphorylation within Syk interdomain A. Analytical modeling shows that multivalent interactions lead to increased WT Syk *cis*-oriented binding by three orders of magnitude. Molecular dynamics (MD) simulations show increased inter-SH2 flexibility in a Y130E phosphomimetic form of Syk, associated with reduced overall helicity of interdomain A. Hybrid MD/worm-like chain polymer models show that the Y130E substitution reduces *cis* binding of Syk. We report computational models and estimates of relative binding for all possible *cis* and *trans* 2:2 Syk:FcεRIγ complexes. Calcium imaging experiments confirm model predictions that *cis* binding of WT Syk is strongly preferred for efficient signaling, while *trans* conformations trigger weak but measurable responses.

## INTRODUCTION

The spleen tyrosine kinase (Syk) is essential for BCR, FcγR and FcεRI signaling, all of which are members of the multichain immunorecognition receptors (MIRRs) family (Johnson et al., 1995; Tamir and Cambier, 1998). Syk activation by MIRRs switches on multiple pathways in mast cells and B cells (Geahlen, 2009; Mocsai et al., 2010), including protein kinase C (PKC) signaling (Kawakami et al., 2000), the PI3K-mediated Akt pathway (Jiang et al., 2003), and transcriptional regulation via the MAPK cascade (Wan et al., 1996). The coupling of Syk to MIRRs relies on a tandem pair of Src homology 2 (SH2) domains in the N-terminal region (Geahlen and Burg, 1994). Both SH2 domains adopt the canonical SH2 fold, comprising a β-sheet flanked by two α-helices (Kuriyan and Cowburn, 1993), and both domains bind to short peptide motifs carrying phosphotyrosine (pTyr) residues (Pawson, 1995). The two SH2 domains are connected by a 45-residue linker region referred to as interdomain A, which consists of three α-helices in a coiled coil conformation and leads to a Y-shaped structure composed of the linker and tandem SH domains (Fütterer et al., 1998). The only other member of the Syk family of non-receptor protein tyrosine kinases is Zap70, which is expressed in T cells and natural killer cells (Chan et al., 1992; Chu et al., 1998; Wang et al., 2010) and whose N-terminal region also adopts a Y-shaped structure (Hatada et al., 1995). In addition to binding pTyr motifs via its tandem SH2 domains (Kihara and Siraganian, 1994), the N-terminal region of Syk/Zap70 is also involved in the autoinhibition of its kinase domain (KD) through residue-residue interactions between interdomain A and the C lobe of the KD (Deindl et al., 2007; Yan et al., 2013). Syk and Zap70 can be functionally homologous for TCR and BCR signaling (Kong et al., 1995; Cheng et al., 1997), although Zap70 is more dependent on Src-family kinases (*e.g.,* Lck) for its catalytic activation compared to Syk (Iwashima et al., 1994; Fasbender et al., 2017).

A distinguishing feature of the MIRRs, which lack intrinsic kinase activity, is the presence of pTyr-containing immunoreceptor tyrosine-based activation motifs (ITAMs) within the cytoplasmic tails of signaling subunits (Reth, 1989; Cambier, 1995; Sigalov, 2005; Harwood and Batista, 2008; Rivera et al., 2008; Smith-Garvin et al., 2009). MIRRs typically incorporate multiple ITAMS, often in disulfide-linked pairs such as the γ homodimers (common to FcγRIII, FcγRI, FcεRI), ζ homodimers (TCR) and Igα,Igβ heterodimers (BCR) (Reth, 2001). We focus here on the high-affinity IgE receptor (FcεRI), which is activated when multivalent ligands crosslink IgE-FcεRI complexes for increased defense against pathogens, wound healing, and allergic responses (Siraganian, 2003; Molfetta et al., 2007; MacGlashan, 2008; Mukai et al., 2018). FcεRI is an αβγ2 oligomeric complex with 3 ITAMs: one in the β subunit and one each in the disulfide-linked γ chains (Blank et al., 1989; Ra et al., 1989). In most models of ITAM-based signaling, phosphorylation of both tyrosines in the same ITAM is assumed to be required for docking of Syk’s tandem SH2 domains (Kihara and Siraganian, 1994; Shiue et al., 1995; Faeder et al., 2003). This *cis* orientation for binding is supported by crystal structures of Syk and Zap70 bound to a single dually tyrosine-phosphorylated peptide (Hatada et al., 1995; Fütterer et al., 1998). Mass spectrometry-based analysis of FcεRIγ found that phosphorylation of the N-terminal Tyr is more abundant than at the C-terminal Tyr (Yamashita et al., 2008), suggesting that ITAM phosphorylation is often incomplete and could limit the availability for *cis*-oriented binding. The common presence of disulfide-linked ITAM-bearing pairs in the MIRRs, as mentioned above, raises the important question: do Syk (and Zap-70) bypass the requirement for full ITAM phosphorylation by docking in *trans* to pairs of pTyrs on adjacent signaling subunits? If so, is this alternative docking orientation equally efficient for signal propagation? We address these questions here through both modeling and experimental approaches.

Given that Syk has two distinct SH2 domains, we consider the evidence that individual SH2 domains show selectivity when interacting with phosphopeptides from their biologically relevant protein binding partners. Prior studies have shown that SH2 domains exhibit 50-1000 fold higher affinity to phosphopeptides derived from binding partners, compared to those with randomized sequences (Panayotou et al., 1993; Songyang et al., 1993; Ladbury et al., 1995). Binding is also enhanced by multivalent interactions (Ottinger et al., 1998). In the FcεRI case, the Syk tandem SH2 domain is bivalent and there are four available binding sites in a fully phosphorylated γ chain homodimer. Other examples of signaling proteins incorporating tandem SH2 domains include the protein tyrosine phosphatases PTPN6 (or SHP-1) (Pei et al., 1996) and PTPN11 (or SHP-2) (Pluskey et al., 1995), as well as phospholipase C-γ1 (Ji et al., 1999). Interestingly, the recruitment of proteins bearing only a single SH2 domain, such as the phosphatase INPP5D (SHIP-1), is thought to be favored by monophosphorylation of BCR ITAMs (O’Neill et al., 2011; Getahun et al., 2016). In the case of Syk, experimentally measured binding affinities indicate micromolar affinity of individual SH2 domains to FcεRIγ while engagement of the tandem SH2 domains results in stronger binding by around three orders of magnitude (Chen et al., 1996). However, the relationship between multivalent binding on Syk:FcεRIγ stoichiometry and signal initiation remains of keen interest.

The ability of tandem SH2 domains to enhance Syk-ITAM occupancy can be impacted by post-translational modifications in interdomain A. We specially focus on Syk recruitment after phosphorylation at site Y130 in interdomain A (Keshvara et al., 1998), since Y130E substitution in Syk resulted in a phosphomimetic recombinant protein with reduced binding of its tandem SH2 domains to phosphorylated ITAMs in co-immunoprecipitation studies (Keshvara et al., 1997; Zhang et al., 2008). This was initially attributed to the structural destabilization of interdomain A and the resulting partial decoupling of both SH2 domains (Zhang et al., 2008; Feng and Post, 2016; Roy et al., 2016). However, high resolution imaging studies in Syk-deficient cells reconstituted with Syk Y130E recombinant protein revealed two interesting observations: 1) Syk recruitment to FcεRI aggregates is retained even for Y130E phosphomimetic, and 2) impaired downstream signaling correlates with an increased off-rate for FcεRI-Syk-Y130E interactions (Schwartz et al., 2017). These results motivate the current studies to understand how interdomain A phosphorylation influences the conformation, binding orientation, and residency time of Syk’s dual SH2 domains on ITAMs.

In this work, we examine Syk-FcεRI binding modes using a combination of conventional and enhanced sampling molecular dynamics (MD) simulations, as well as hybrid MD/polymer theory (Sethi et al., 2011). We first developed an analytical model based on simple structural arguments to evaluate how multivalent interactions cause a three orders of magnitude increase in binding compared to individual SH2 domains. Next, conventional MD simulations show that the Y130E substitution leads to increased flexibility in the relative orientations of the two SH2 domains in Syk. Replica exchange MD (REMD) simulations of interdomain A further show that the amount of helical structure in this linker region is reduced by Y130E, impacting the distance between the tandem SH2 domains. We then show using a hybrid approach of MD simulations and worm-like chain (WLC) polymer models that the combination of higher inter-SH2 distance and increased flexibility in the Syk Y130E mutant reduces its binding affinity to FcεRIγ in a *cis* orientation. We also present structural models and binding energy estimates for the multiple *trans* binding modes that are possible for binding of two wildtype (WT) Syk molecules (one pair of tandem SH2 domains each) to a dimer of phosphorylated FcεRIγ chains (one pair of pTyr motifs each). Overall, the Syk tandem SH2 domains binds best in the *cis* orientation, followed by a diagonally opposed *trans* orientation targeting the N pTyr in one ITAM and the C pTyr in the other ITAM. Predictions of the computational modeling were supported by experiments in rat basophilic leukemia (RBL-2H3) cells expressing endogenous WT Syk and transiently expressing chimeric receptors bearing the FcεRIγ cytoplasmic tail with either WT or mutant ITAM sequences.

## RESULTS

### Multivalent binding of wildtype Syk tandem SH2 domains to a single FcεRIγ chain

We first constructed a structure-based analytical model that describes how local concentration effects lead to an enhancement of overall *cis* binding between the linked SH2 domains from Syk and the paired pTyrs in a FcεRIγ chain ITAM. In **Fig. 1A**, the initial binding steps of either N-SH2:C-ITAM or C-SH2:N-ITAM are shown with equilibrium association constants of *K_N_* or *K_c_*, respectively. The subsequent binding steps involve either i) binding of C-SH2:N-ITAM after N-SH2:C-ITAM with an effective equilibrium association constant of
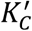, or ii) binding of N-SH2:C-ITAM after C-SH2:N-ITAM with an effective equilibrium association constant of
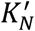.
Both
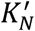 and
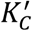 are given by multiplying *K_N_* and *K_c_* by a factor *C_eff_* that represents the effective concentration of unbound ITAM that unbound SH2 experiences upon binding of the other SH2:ITAM pair. This effective concentration factor is the same in both pathways described above (either binding of N-SH2:C-ITAM first or binding of C-SH2:N-ITAM first). This model is similar to the one we used previously for investigating the Grb2-Sos1 multivalent complex (Sethi et al., 2011), where the effective equilibrium association constant for binding of both motifs, *K_overall_* (red arrows in **Fig. 1A**), is given by

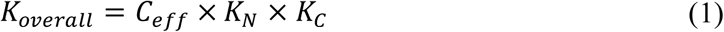

or the product of both monomeric association constants with the effective concentration.

**Figure 1.**
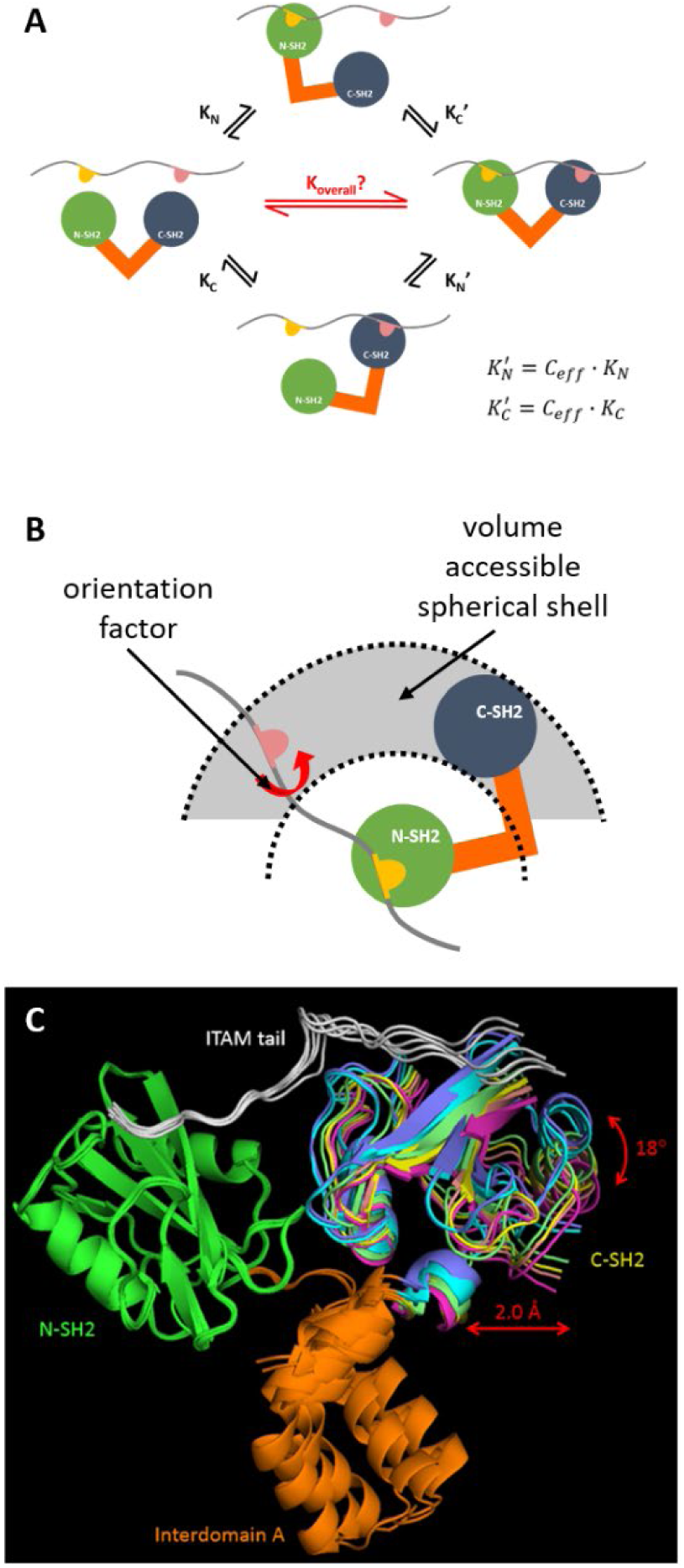
Structure-based analytical model for multivalent binding of Syk tandem SH2 domains to a single FcεRIγ chain. (A) Reaction network from unbound (left) to dual-bound (right) Syk. The N-SH2, C-SH2, and interdomain A are shown in green blue, and orange, respectively. The unstructured γ chain is shown as a gray wavy line, with pY64 and pY75 colored pink and yellow, respectively. (B) Schematic showing the accessible spherical shell where unbound SH2 can search for unbound pTyr given prior binding of the other SH2:pTyr pair, and an orientation factor representing the flexibility of the γ chain. (C) The six complexes in the asymmetric unit of the crystal structure of Syk tandem SH2 domains bound to CD3 s (PDB 1A81) (Fütterer et al., 1998) were structurally aligned at the N-terminal SH2 (green cartoons). The translational (2 Å) and orientational (18°) variability range for C-terminal SH2 (magenta to blue cartoons) among these six aligned complexes are shown with red arrows.

To derive a relation for the effective concentration, we modeled the second binding step using the top path in **Fig. 1A**, although the other direction would give the same results. The binding of the first SH2 domain restricts the region in space that the second SH2 domain can access. The effective concentration of sites the second domain can bind to is just the number of unbound pTyr in the accessible region that are oriented such that binding can occur, divided by the volume of the accessible region *V_acc_*. We model *V_acc_* as a spherical shell (colored gray in **Fig. 1B**) with the bound SH2:ITAM pair at the center of the sphere. Based on the crystal structure of Syk tandem SH2 domains bound to CD3ε (PDB 1A81) (Fütterer et al., 1998) that contained six complex structures in the asymmetric unit cell, the spherical shell is located between 3.2 nm to 3.4 nm from the sphere center (C-SH2 translational variability of 2 Å in **Fig. 1C**). The FcεRIγ chain also has an extended backbone in the crystal structure such that the inter-pTyr orientation can be assumed to be uniformly distributed between 0° to 360°, however only a fraction of these orientations are found to lead to successful complexes with Syk (C-SH2 orientational variability of 18° in **Fig. 1C**). This can be expressed as an orientation factor (*f_or_*) of 18°/360° or 0.05, which is the probability of the extended FcεRIγ chain adopting an inter-ITAM orientation that allows multivalent binding with Syk. The effective concentration is thus given by

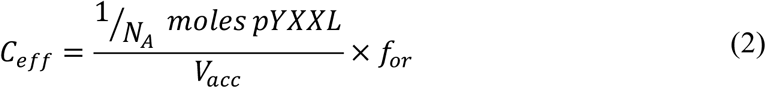

where *N_A_* is the Avogadro constant. We note that the binding orientation found in the crystal structure, where N-SH2 interacts with C-ITAM and C-SH2 with N-ITAM, was the only *cis* mode considered here. This is because a physical atomistic model cannot be built for the other *cis* mode, with interactions between N-SH2:N-ITAM and C-SH2:C-ITAM, since the linker sequence between both ITAMs on a single γ chain is not long enough to accommodate these interactions (**Supp. Fig. 1**).

For the system shown in **Fig. 1C**, the estimated *C_eff_* is around 3.03 mM. The experimentally-measured equilibrium dissociation constants for binding of doubly-phosphorylated FcεRIγ tail to N-SH2 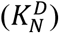 and C-SH2 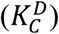 are >2.3 μM and 1.3 μM (Chen et al., 1996), respectively, whose reciprocals are *K_N_* and *K_c_*. Plugging these values into equation (1) and getting the reciprocal of *K_overall_* gives an estimated value for the effective equilibrium dissociation constant 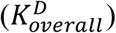 of around 1 nM, which provides only a lower bound since the value used for 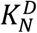 is also only a lower bound. The experimentally-measured value is 1.4 nM (Chen et al., 1996). Thus, this simple analytical model captures the three orders of magnitude multivalent effect observed when going from monovalent binding (with μM affinity) to bivalent binding (with nM affinity) between WT Syk tandem SH2 domains and the two pTyrs on a single FcεRIγ chain.

### Changes in inter-SH2 distances in models of the Syk Y130E phosphomimetic

Our next goal was to apply unbiased MD simulations to investigate the impact of interdomain A phosphorylation on the *cis* binding of Syk’s tandem SH2 domains, using the Y130E phosphomimetic as a model system. The challenge of applying the earlier analytical model for Syk Y130E mutant is that there is no experimentally-resolved structure for this mutant. We thus turned to MD simulations to investigate how the Y130E mutation affects the structure of the tandem SH2 domains. We applied two order parameters to quantify the flexibility between SH2 domains in our simulations (**Fig. 2A**). The first coordinate was the distance between Cα’s of R21 on N-SH2 and R174 on C-SH2 that are both part of the pTyr-binding sites on their respective SH2 domains, and the second coordinate was the dihedral angle involving the Cα’s of R21 and L28 on N-SH2 with R174 and V181 on C-SH2. These coordinates provided measures of the inter-SH2 distance and orientation, respectively. For both constructs, seven simulation replicates were performed for 2 μs simulation time per replicate.

**Figure 2.**
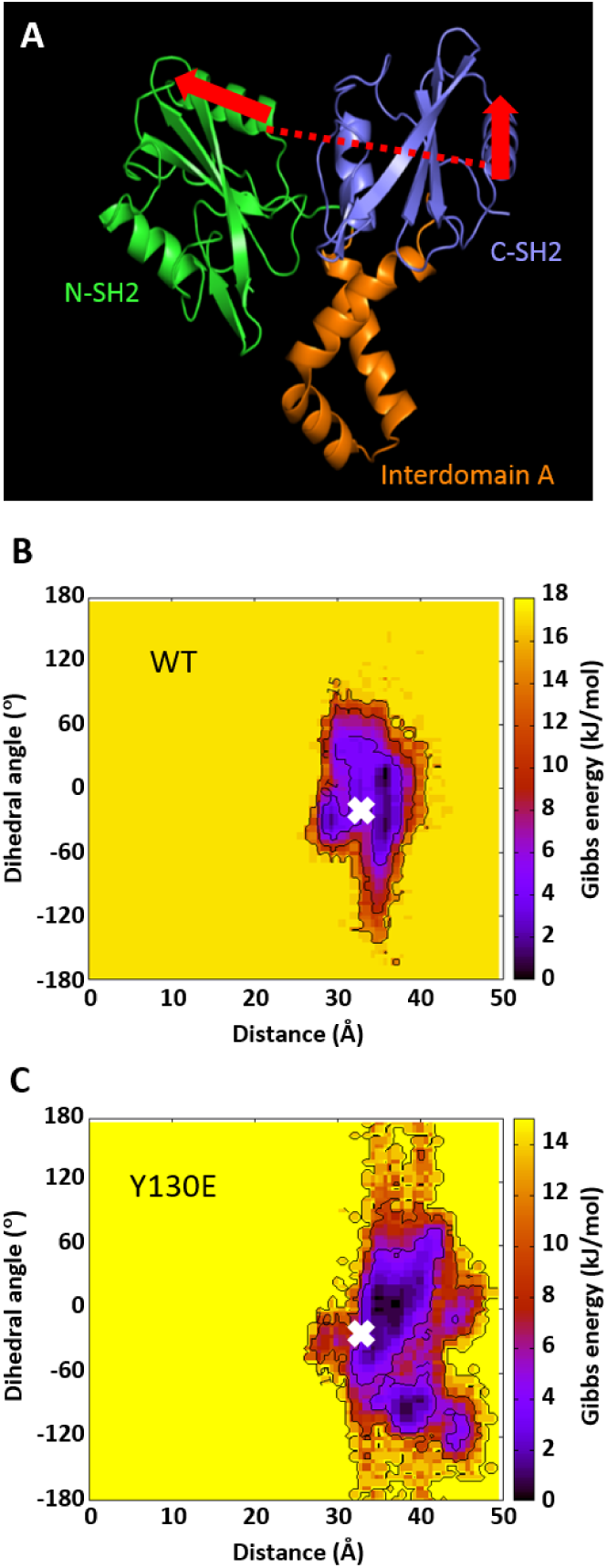
Unbiased MD simulations show higher inter-SH2 flexibility in the Y130E phosphomimetic form of Syk. (A) Distance (red dashed line) and dihedral (across red arrows) reaction coordinates for describing the inter-SH2 positions and orientations. (B) Free energy surface map for the WT Syk simulations based on the two reaction coordinates. (C) Corresponding free energy surface map for the Y130E phosphomimetic Syk. The white X in (B) and (C) gives the position of the starting conformation for the MD simulations. Data was collected from seven replicates per construct, with 2 μs simulation time per replicate.

These runs showed that the Y130E phosphomimetic has higher translational and orientational flexibility than WT Syk (**Fig. 2 B and C**). We then chose two replicates per construct and extended the simulation time to 6 μs per replicate, and found the same higher flexibility trends for the mutant compared to WT (**Supp. Fig. 2 A and B**). The higher flexibility seen in the simulations of the WT construct (**Fig. 2B**) compared to the crystal structure (**Fig. 1C**) is because the latter had a bound dually-phosphorylated ITAM tail that likely constrained the inter-SH2 flexibility. Closer inspection of these simulations showed that, in contrast with experimental findings, there was no destabilization of the helical structure of interdomain A in the Y130E phosphomimetic (**Supp. Fig. 3**). We instead observed changes in the number of domain-domain contacts for Syk due to Y130E, particularly a decreased number of contacts between N-SH2/C-SH2 and C-SH2/interdomain A as well as an increased number of contacts between N-SH2/interdomain A (**Supp. Fig. 4**); these features likely account for the increased flexibility of the Y130E mutant. We note that verification of the structural destabilization of interdomain A is not accessible within the time scales of unbiased MD simulations considered here.

To address this, we next used an enhanced sampling simulation technique called replica exchange MD (REMD) that allows more efficient sampling of the conformational ensemble for the system under study (Sugita and Okamoto, 1999; Garcia and Sanbonmatsu, 2001). The REMD simulations were limited here to residues K115 to T159 that comprise interdomain A. The simplification by removal of both SH2 domains allows for a smaller system size to enhance the conformational sampling of the interdomain. For the WT and Y130E constructs, we ran 63 replicas each within the temperature range 275-475 K, with 1.2 μs of simulation time per replica for a cumulative simulation time of 75.6 μs. Analysis was performed for each construct cumulatively on the fifteen replicas with temperatures less than 310 K.

We again counted the number of helical bonds as a measure of the stability of the helical structure of interdomain A for both constructs in these simulations. Overall, we found that the Y130E mutant showed more helical unfolding than the WT construct, with around five helical bonds lost for Y130E compared to around two helical bonds for WT by the end of the simulations (**Fig. 3A**). For the three individual helices that comprise the interdomain A structure, we found that more unfolding occurred in helix 1 that contains the Y130 phosphorylation site, with the mutant destabilizing around three helical bonds compared to < 1 for WT (**Fig. 3B**). In contrast, only around one helical bond was destabilized in helices 2 and 3 for the phosphomimetic mutant (**Fig. 3 C and D**).

**Figure 3.**
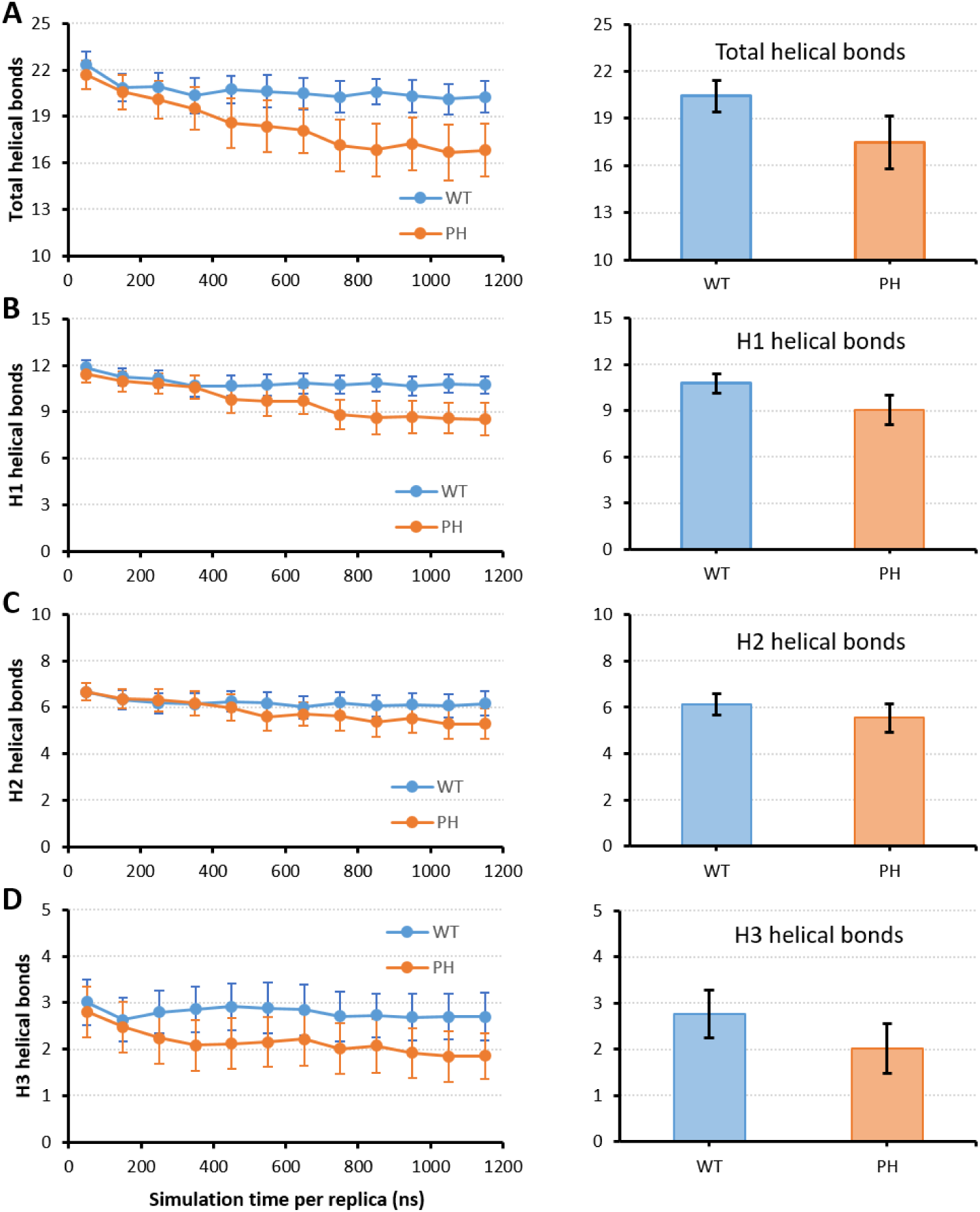
REMD simulations show more helical unfolding of interdomain A in the Y130E phosphomimetic mutant. Time profiles over the entire trajectory (left) and cumulative bar plots of the last 500 ns (right) for the number of helical bonds from REMD simulations of WT (blue) and Y130E mutant (orange) interdomain A. Helical bond counts were done for (A) total helices, (B) helix 1 (H1), (C) helix 2 (H2), or (D) helix 3 (H3). Error bars give s.e.m.

Given the helical unfolding observed here for Y130E, we were next interested to see how this may affect the distances between the SH2 domains. Thus, for the conformational ensembles obtained for the WT and Y130E interdomain A constructs in these REMD simulations, we reattached both SH2 domains *in silico* to every snapshot in order to assess the inter-SH2 distance distributions for both constructs. Given that the two ends of interdomain A were free to move in the REMD simulations, we filtered for only those models containing interdomain A and reattached SH2 domains that did not show steric clashes between N-SH2, C-SH2, and interdomain A. About 5% from the total number of models in both constructs remained physically viable after performing this filtering.

We found that the distance distributions for both constructs were broader from the REMD-based models compared to those from the unbiased simulations (**Fig. 4 A and B**; compare with x axes in **Fig. 2 B and C**), but consistently showed that the Y130E mutant samples more inter-SH2 distance values than WT. The mean inter-SH2 distance values for the WT construct were very similar at around 32 Å between the REMD-based models and the unbiased simulations. For the Y130E construct, however, the mean inter-SH2 distance was larger for the REMD-based models (around 46 Å) compared with the unbiased simulations (around 36 Å), which is consistent with the increased structural instability of interdomain A seen in the REMD simulations. Using these distance distributions, we can employ a hybrid MD/polymer approach to estimate the multivalent effect on binding for WT and Y130E mutant Syk to FcεRI.

**Figure 4.**
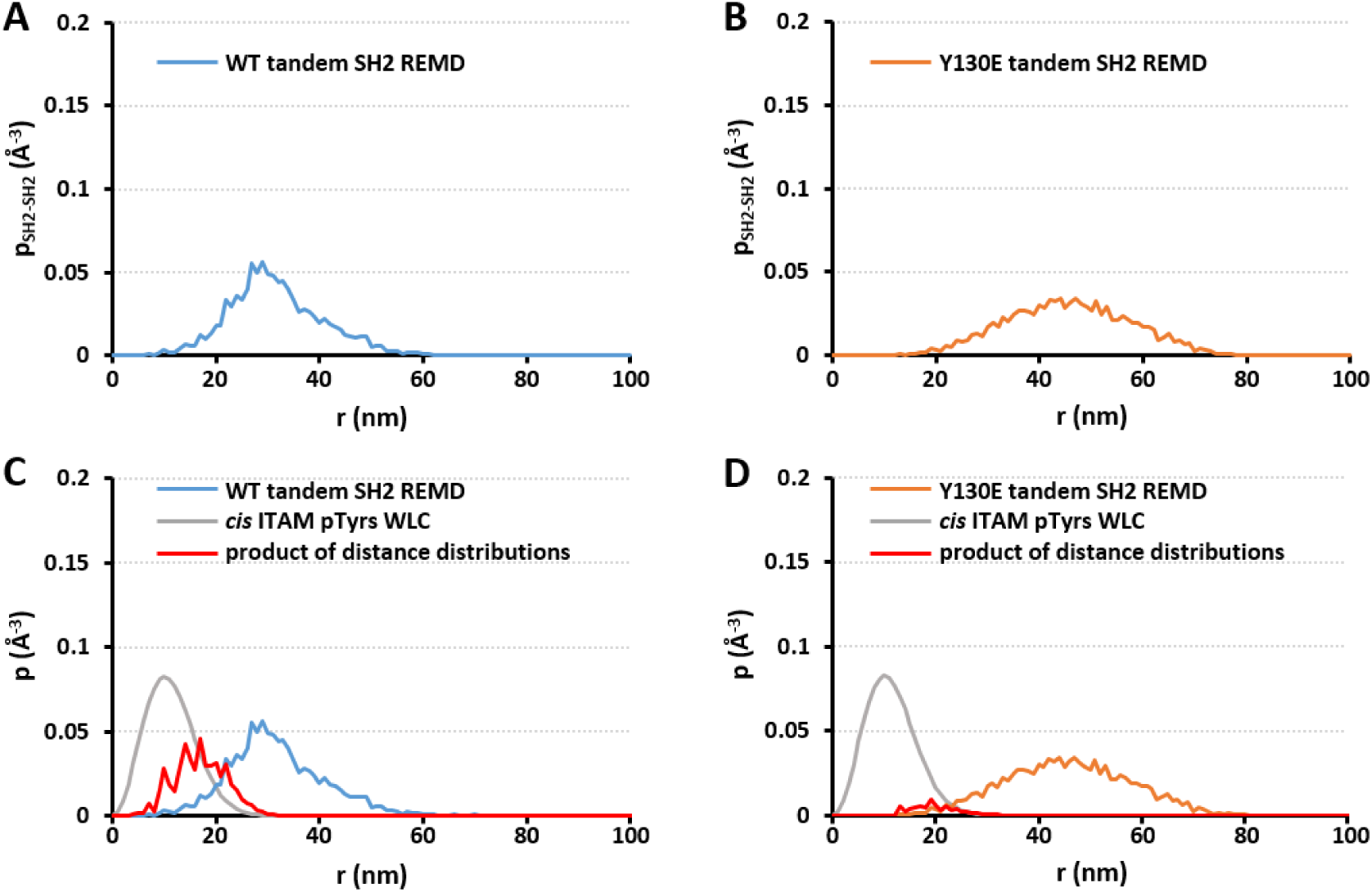
Distance distributions between tandem SH2 domains from REMD-based models and between ITAM pTyrs from WLC models. (A) Distribution between SH2 domains from REMD-based models of WT tandem SH2. (B) Distribution between SH2 domains from REMD-based models of Y130E mutant tandem SH2. (C) The WT Syk distribution from (A) is shown with the distribution between ITAM pTyrs from the WLC model of a single FcεRIγ chain for *cis* binding (gray curve). (D) The Y130E Syk distribution from (B) is shown with the distribution between ITAM pTyrs from the WLC model of a single FcεRIγ chain for *cis* binding (gray curve). In (C-D), the product of both distance distributions is shown by the red curve, whose integrated area gives *C_eff_*. The magnitudes of the red curves have been increased here by 100X to facilitate their visualization and comparison.

### Hybrid MD/polymer model for estimating multivalent binding of Y130E phosphomimetic Syk to a single FcεRIγ chain

As mentioned earlier, multivalent binding for Syk:FcεRI depends on the effective concentration *C_eff_* of unbound ITAM that unbound SH2 experiences when the other SH2:ITAM pair is bound. Given distance distributions between the Syk SH2 domains (*𝑝_SH2–SH2_*) and between the ITAM pTyrs (*𝑝_ITAM–ITAM_*), it has been shown that *C_eff_* can be calculated by determining the overlapping regions between both distributions and integrating over the products of the probabilities in these regions for all possible distances (Van Valen et al., 2009; Sethi et al., 2011):

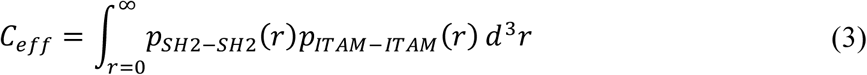

Here, we use the distance distributions derived from the REMD-based models of WT and Y130E mutant Syk tandem SH2 domains as the corresponding *𝑝_SH2–SH2_* in equation (3). To derive *𝑝_ITAM–ITAM_*, we treat the FcεRIγ chains using a WLC polymer model. For binding of a Syk tandem SH2 to a single FcεRIγ chain (*i.e., cis* binding):

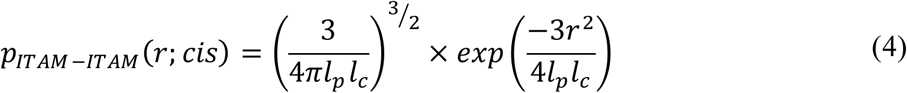

where *l_p_* and *l_c_* are the persistence and contour lengths of the peptide between the two pTyrs in the single FcεRIγ chain. Both of these length values can be expressed as a function of the number of residues in the peptide. The distance distribution plot for the *cis* binding WLC model is shown overlapped with corresponding plots for the WT and Y130E tandem SH2 REMD-based models in **Fig. 4 C and D**. The *C_eff_* values (integrated area under red curves in **Fig. 4 C and D**) for binding of a single FcεRIγ chain to WT and Y130E mutant Syk are estimated at 7.9 mM and 1.1 mM, respectively. These correspond to 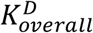 values of 0.38 nM and 2.8 nM, respectively, indicating that WT Syk has stronger affinity than the Y130E mutant for *cis* binding to a single FcεRIγ chain by around an order of magnitude.

### *Trans*-oriented binding modes increase the complexity of Syk-FcεRI multivalent interactions

The above hybrid MD/polymer model assumes that Syk only binds via a *cis* mode to an FcεRIγ monomer. However, an outstanding question is whether Syk can also utilize *trans* binding modes that span pTyr residues in both chains of an FcεRIγ dimer. We applied computational modeling to explore how Syk molecules can bind to the ITAMs in the FcεRIγ dimer, using a maximum stoichiometry of two Syk molecules (total of four SH2 domains) to one FcεRIγ dimer (total of four pTyrs). We assume here that all ITAM tyrosines have been phosphorylated. From a total of 24 possible permutations, we found that only six of these lead to viable atomistic models after filtering out structurally redundant or physically unrealistic models. One of these six models comprises the *cis* binding mode while the other five models comprise different *trans* binding modes (**Fig. 5A**). Three of the five *trans* binding modes contain variations of *trans* (NC) binding (Trans1, Trans2, and Trans3 in **Fig. 5A**), while the remaining two (Trans4 and Trans5) combine *trans* (NN) and *trans* (CC) binding.

**Figure 5.**
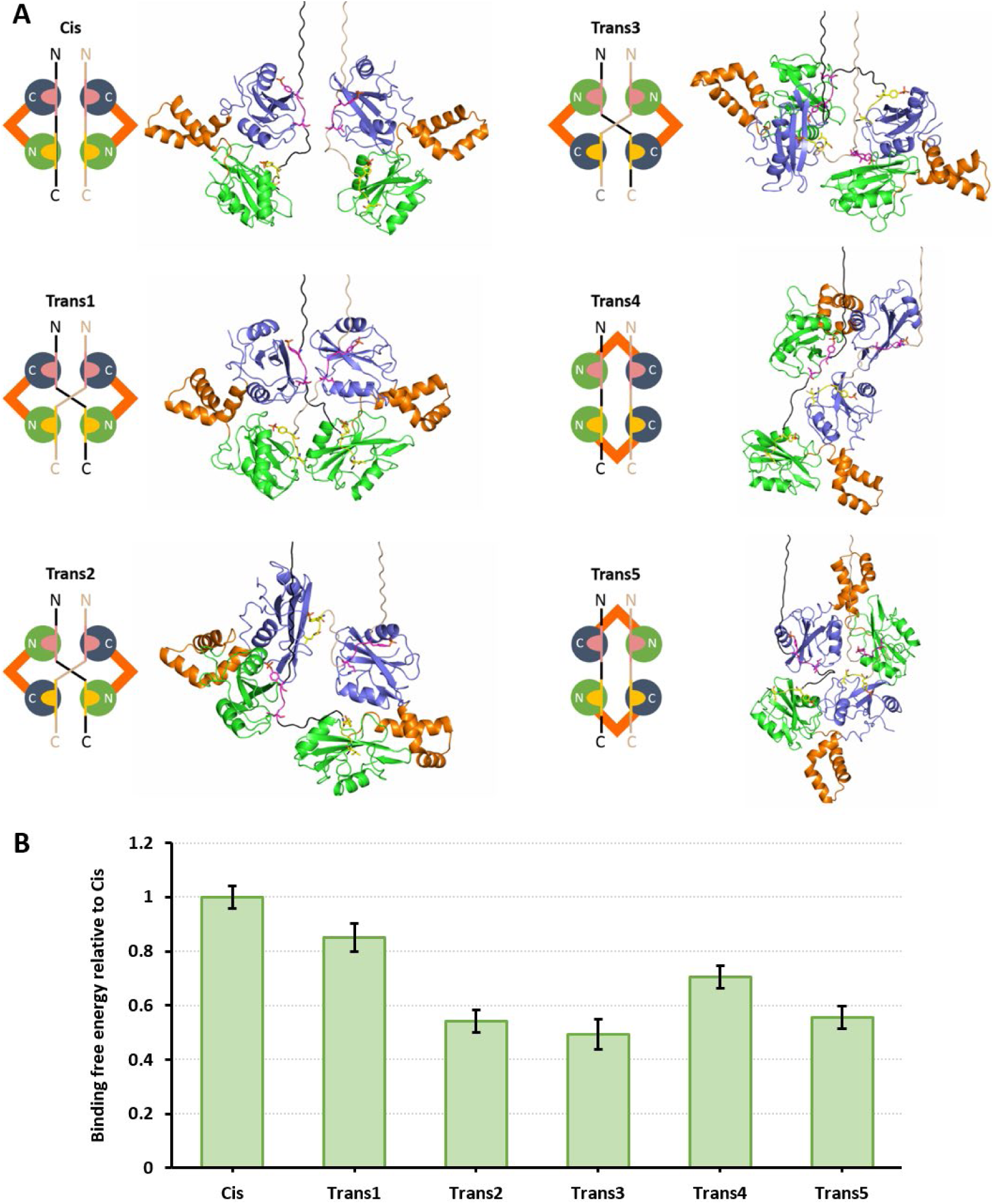
WT tandem SH2 domains show higher affinity for *cis* binding in complex models of Syk:FcεRI with 2:2 binding stoichiometry. (A) Diagrams and corresponding atomistic structures of one *cis* and five *trans*binding modes that are physically viable and consistent with a 2:2 binding stoichiometry. Syk is depicted here using cartoons for N-SH2 (green), C-SH2 (blue), and interdomain A (orange). In each model, the two γ chains are colored differently (black and light pink), and showing how pY64 (pink) and pY75 (yellow) are bound. Portions of the linker that connect the N-terminal end of the ITAM to the γ chain transmembrane helices are shown for each model using artificial extended conformations, to give a sense of how each model may be oriented relative to the surface of the cytoplasmic leaflet of the cell membrane. (B) Unbiased MD simulations of these models in solution were then performed, with the artificial linker regions that connect the ITAMs to the transmembrane helices removed. The overall binding free energies were then calculated using the linear interaction energy (LIE) method (Aqvist et al., 1994) and plotted here relative to the *cis* binding free energy. Error bars give s.e.m.

Unbiased MD simulations for each of these 2:2 models of Syk:FcεRIγ were then performed using the soluble portions of the complexes, with three replicates per model at 1 μs simulation time per replicate. For evaluating the relative stability of each of these models, we used the linear interaction energy (LIE) method (Aqvist et al., 1994) to compute the total binding free energies of the ITAMs to their corresponding SH2 domains in each model. Results show that the *cis* orientation results in the strongest binding free energy for WT Syk at a 2:2 stoichiometry between Syk:FcεRIγ (**Fig. 5B**). Among the *trans* modes, Trans1 binding is strongest (80% of *cis*) followed by Trans4 (~70% of *cis*). Trans 2, 3 and 5 binding modes were feasible, but with interaction strengths of approximately half that of *cis* binding.

We first assessed whether these differences in stability arise from interactions of individual Syk SH2 domains with the individual γ chain pTyrs: the N-terminal pTyr site (pY64) or the C-terminal pTyr site (pY75). As shown in **Supp. Fig. 5**A, we computed interaction energies for each pair and found similar results for each ITAM:SH2 pair regardless of the binding mode. Exceptions may be pY75:N-SH2 in the Trans2 and pY64:N-SH2 binding in the Trans5 orientations, since unbinding occurred for each in one out of three simulation replicates. Thus, the binding modes do not appear to strongly influence individual Syk SH2 interactions with ITAM pTyrs.

It is noteworthy that the interaction energies for binding of Syk’s C-SH2 domain to pY64 in all orientations are predicted to be slightly stronger than binding of the same C-SH2 domain to pY75 (**Supp. Fig. 5B**). Conversely, the binding of Syk’s N-SH2 domain to pY75 is predicted to be slightly stronger than its binding to pY64. **Supp. Fig. 5B** also shows that, on average, Syk C-SH2 interactions with either pTyr site are stronger than the corresponding N-SH2 interactions. For most binding orientations, the simulations suggest that the interactions by Syk’s N-SH2 domain are comparable. These results are qualitatively similar to experimental measurements of binding constants between individual Syk SH2 domains and CD3ε ITAMs, in the order pY64:C-SH2 > pY75:C-SH2 > pY75:N-SH2 > pY75:N-SH2 (Feng and Post, 2016).

We next considered the possibility that the engagement of tandem SH2 domains with both pTyrs of the ITAM induces additional contact sites. As shown in **Fig. 6A**, we made the novel observation that the interaction stability of a Syk tandem SH2 domain bound to a dually phosphorylated ITAM is improved by additional residue-residue interactions (red circle) of the C-SH2 domain with pY75 (that is bound to N-SH2). These interactions likely contribute to the stronger binding observed in Cis and Trans1 orientations (**Fig. 6B**). Note that these unexpected contacts are asymmetric, as pY64 (that is bound to C-SH2) is not capable of making additional interactions with N-SH2.

**Figure 6.**
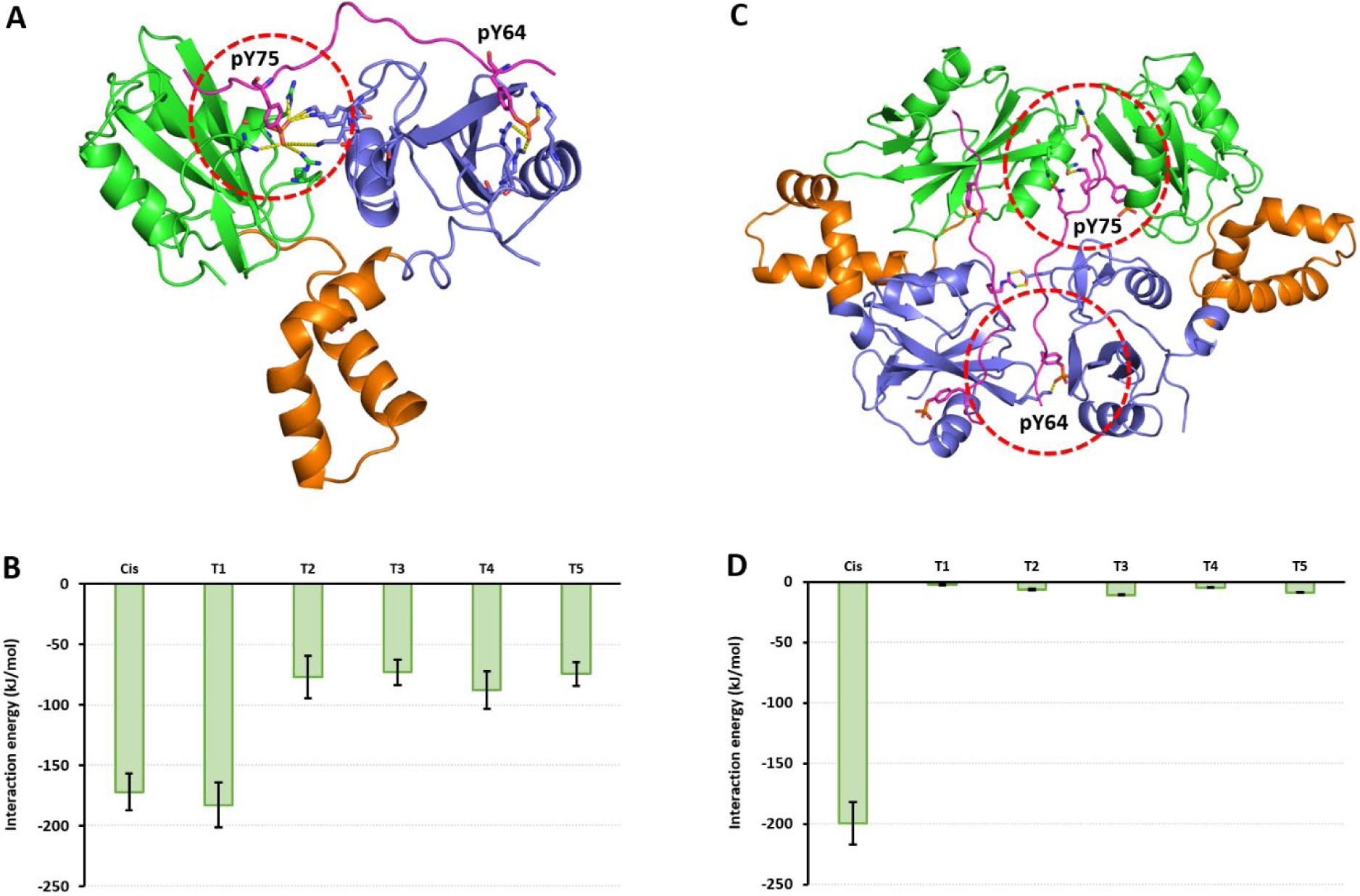
Additional contacts contribute to further stabilization of the *cis* binding mode. A) Illustration showing that in the binding of Syk N-SH2 domain to pY75 and C-SH2 domain to pY64, asymmetrical contacts between pY75 and C-SH2 are formed (red encircled region). This is not the case for pY64 and N-SH2. Key: N-SH2 (green), C-SH2 (blue), interdomain A (orange), γ chain (magenta). (B) Average interaction energies of these asymmetrical contacts when engaged in each of the different 2:2 binding modes. Error bars give s.e.m. C) Illustration of additional contacts between the pTyrs bound to the Syk molecule on the right with the “bystander” Syk molecule on the left (red circles) for the *cis* orientation of the 2:2 SykFcεRIγ complex. Proteins are colored as in (A). (D) Average interaction energies of these additional contacts, illustrating that these are significant only in the *cis* orientation. Error bars give s.e.m.

**Fig. 6C** indicates that additional asymmetrical contacts further contribute to the stability of Syk docking when engaged in 2:2 Syk:FcεRIγ complexes. The structural model shows that the pTyrs and nearby residues of the γ chain bound to the Syk molecule on the right engage in residue-residue contacts with the Syk molecule on the left (red circles). Since these interactions likely are unique to the *cis* orientation (**Fig. 6D**), these data provide mechanistic support for *cis*-binding mode as the most stable form of multivalent binding between Syk and FcεRIγ (**Fig. 5B**).

### *Trans* binding modes generate weak but measurable signaling in live cells

Experimental measures of calcium mobilization provide a sensitive indicator of FcεRI and Syk activation in cultured mast cells (Smith et al., 2001). We used an established model system to study the individual contribution of the FcεRIγ subunit in rat basophilic leukemia (RBL-2H3) cells, based upon expression of a chimeric TT-γ receptor derived from the coding sequences of extracellular and transmembrane domains of the Tac antigen (IL2α subunit) fused in frame with the γ subunit cytoplasmic tail sequence (Letourneur and Klausner, 1991; Wilson et al., 1995). For this study, we prepared transient expression vectors to express either WT chimeric receptors (TT-γ WT) or mutant versions with tyrosine to alanine substitutions in each of the two ITAM tyrosines (TT-γ Y64A and TT-γ Y75A). The chimeric receptors are first converted to a dimer state by pretreating cells with a monoclonal anti-Tac antibody, consistent with the γ homodimer state in the resting FcεRI complex. This pretreatment does not initiate significant changes in cytosolic calcium (**Supp. Fig. 6A-E**). Syk-dependent signaling leading to calcium mobilization is then initiated by addition of polyclonal anti-Tac antibodies that aggregate TT-γ dimers (Wilson et al., 1995). Results of this model system are reported in **Fig. 7**, based upon ratio imaging of cells loaded with the calcium reporter, Fura-2AM (Smith et al., 2001). As expected, antigen-mediated crosslinking of intact, IgE-primed FcεRI complexes led to marked elevation of cytoplasmic calcium in 95% of cells (**Fig. 7E** and **Supp. Fig. 7A)**. Aggregation of TT-γ WT chimeric receptors led to robust calcium response in 41% of the cells examined, with weak, oscillatory calcium bursts in an additional 2% of cells (**Fig. 7 A and E**). Measurable responses were observed after aggregation of either of the TT-γ mutant forms, where only monophosphorylation is possible (TT-γ Y64A and TT-γ Y75A; **Fig. 7 B and C**). However, as reported in **Fig. 7E**, these responses were limited to a small fraction of cells examined (4-7%). The profiles of these responses typically exhibited a delayed onset, a lower overall magnitude of change, or were oscillatory (see **Supp. Fig. 6F** for representative calcium traces used to score responding cells).

**Figure 7.**
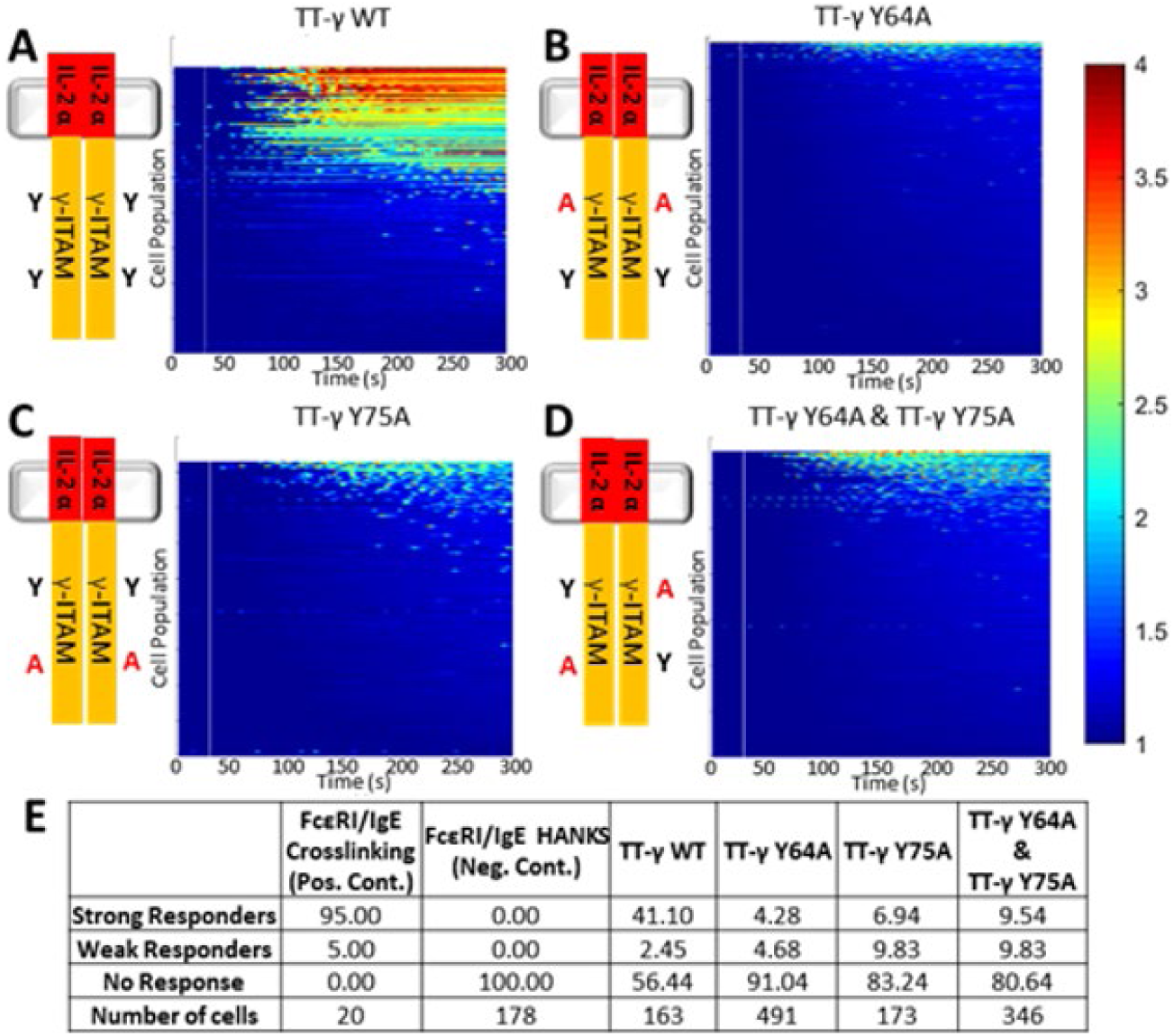
Ca^2+^ imaging shows Syk(WT) SH2 domain preference for *cis* but existence of *trans*initiated signaling. Calcium imaging signaling responses resulting from (A) TT-γ WT, (B) TT-γ Y64A, (C) TT-γ Y75A, and (D) co-expressed TT-γ Y64A and TT-γ Y75A. Heatmaps indicate Ca^2>+^ release, with red indicating a strong response and blue indicating no release. White vertical line indicates time of secondary antibody addition. (E) Table summarizing the calcium responses for crosslinking of intact FcεRI, negative controls, and the different TT-γ constructs (see also **Supp. Figs. 6** and **7**).

We also co-transfected both TT-γ Y64A and TT-γ Y75A into RBL-2H3 cells and evaluated calcium mobilization after addition of antibodies to produce mixed aggregates containing the two possible monophosphorylated forms of the γ ITAM. Consistent with the highest *trans* binding values reported in **Fig. 5B** for a Trans 1 “diagonal” binding mode, we observed that a slightly higher percentage of cells (>9 %) mounted a strong and persistent calcium response (**Fig. 7D and E**), with another 9% producing weaker responses. These data sets led to the conclusion that Syk recruitment is possible in receptor aggregates that only provide *trans* binding modes but that signaling output is weak compared to aggregates that also permit Syk binding in the canonical *cis* orientation.

## DISCUSSION

Efficient immune cell signaling relies on protein-protein interactions that are often facilitated by reversible phosphorylation events (Hunter, 1995; Pawson and Scott, 2005; Basson, 2012; Ardito et al., 2017; Gelens et al., 2018). The recruitment of the Syk/Zap-70 family of cytosolic tyrosine kinases to ITAM motifs is an essential step in signaling by the MIRR family of immunoreceptors (Geahlen, 2009; Mocsai et al., 2010). In this work, we have applied multiple computational modeling approaches to characterize the structural features that control interactions between Syk and the FcεRI γ-ITAM. We report contributions of multiple factors to the specificity and strength of the Syk-FcεRI complex, including multivalency and *cis* versus *trans* binding orientations. We unveil novel residue-residue interactions made possible only in specific orientations. We also provide mechanistic insight into the impaired binding of the Syk-Y130E that mimics structural changes induced by phosphorylation in interdomain A.

The influence of multivalency was explored through develoμMent of a structure-based analytical model. We show that local concentration effects arise due to the bivalent nature of Syk’s tandem SH2 domains together with the paired pTyrs in the FcεRI γ-ITAM. We applied an effective equilibrium association constant for binding of the linked motifs, previously derived (Sethi et al., 2011), that comprises the product of the two monomeric association constants and a quantity called *C_eff_* as shown in equation (1). *C_eff_* represents the effective local concentration of pTyr available for SH2 binding, given that one SH2:pTyr is already engaged. We used the translational variability between the Syk tandem SH2 domains observed in the six structures from PDB 1A81 (Fütterer et al., 1998) to define the volume of the spherical shell where unbound SH2 can search for unbound pTyr. The orientational variability between the tandem SH2 domains in these six structures was also used to obtain a factor for the fraction of orientations within this search shell that allow for successful binding events. Combining these values with the experimentally-measured monomeric binding constants (Chen et al., 1996), we arrived at a calculated effective binding constant of ~1 nM for the system that is consistent with the experimentally-measured affinity for Syk SH2-SH2 bound to a dually phosphorylated ITAM (Chen et al., 1996).

Binding characteristics of the Syk Y130E mutant are particularly of interest, since expression of this phosphomimetic has impaired signaling capability (Schwartz et al., 2017). Since there is no resolved structure for the phosphomimetic, we used a hybrid MD/polymer model to generate two distance distributions (inter-SH2 for Syk and inter-pTyr for FcεRIγ). First, the inter-SH2 distance distribution was computed from unbiased MD simulations of the Syk tandem SH2 domains connected by the interdomain A linker using WT and Y130E mutant constructs. These simulations showed the mutant sampling conformations that cover a wider distance distribution than the WT (x-axes in **Fig. 2 B and C**). The mutant also shows larger orientational variability in the conformations it sampled in these simulations (y-axes in **Fig. 2 B and C**). These conformational differences were attributed to changes in the number of domain-domain contacts in the mutant relative to the WT construct (**Supp. Fig. 4**). In particular, there is a decrease in the number of contacts between the N-SH2 and C-SH2 domains in the presence of the mutation, which is consistent with the experimentally-observed partial decoupling of both SH2 domains in the mutant (Zhang et al., 2008; Feng and Post, 2016; Roy et al., 2016).

MD simulations provided insufficient information on the influence of the Y130E substitution on helical stability, due to limited sampling even at microsecond time scales. Motivated by experimental observations that Y130E does disrupt helical stability (Zhang et al., 2008; Feng and Post, 2016; Roy et al., 2016), we computed inter-SH2 distance distribution from REMD simulations of the interdomain A linker. Only the linker was included in these simulations in order to enhance the sampling of adopted conformations and to evaluate the impact of the Y130E mutation. The two SH2 domains were then reattached to each conformation in the WT and Y130E interdomain A ensembles, from which the inter-SH2 distance distribution was obtained after filtering of models that showed large steric clashes upon adding back both SH2 domains. The models based on the REMD simulations of Y130E linker showed more unfolding of helical turns compared to the WT model (**Fig. 3**). The inter-SH2 distance distribution for Y130E in these simulations was broader and had a larger average value compared to WT (**Fig. 4 A and B**).

A major uncertainty in these calculations is that the FcεRIγ cytoplasmic tail is structurally disordered; adequate conformational sampling of these unstructured regions is challenging for MD simulations (Lopez et al., 2015) even using enhanced sampling techniques. To evaluate the inter-pTyr distance distribution within an ITAM, we used a WLC polymer model. When applied to protein chains, WLC modeling derives this distance distribution based on equation (4) as a function of the persistence and contour lengths (gray curves in **Fig. 4 C and D**). These length values provide the minimum length where the polymer can be modeled as a random walk and the maximum possible length of the polymer, respectively (Zhou, 2001; Ohashi et al., 2007; Rawat and Biswas, 2009). The product of both distributions at each distance value is shown as a red curve, and integrating the area under this curve gives an estimate of *C_eff_* as given in equation (3). The effective binding constant for the Y130E Syk mutant was found to be approximately 10-fold lower than WT. Taken together, these data support the experimental observation that the Y130E phosphomimetic form of Syk can be recruited to FcεRI complexes. If the 10-fold lower binding is mainly attributable to an increased *k_off_* (*i.e.*, faster dissociation rate of formed complexes), then this would be consistent with the experimentally-observed shorter lifetimes of bound Y130E mutant Syk (Schwartz et al., 2017).

Many MIRRs incorporate paired ITAM-bearing subunits, including the FcεRIγ_2_ subunit (shared with Fcγ receptors), as well as TCRζ_2_ and the BCR Igα-Igβ complex. These paired homodimers and heterodimers are stable by virtue of a disulfide linkage (Turner and Kinet, 1999; Call et al., 2006). This markedly expands the complexity and number of variations for possible multivalent interactions, by comparison to the simplest case considered in **Fig. 1** for Syk tandem SH2 domains bound to paired pTyrs in the *cis* orientation. We used molecular modeling and simulations to explore the possibility that Syk can also stably bind to γ homodimers via several *trans* orientations. Simulations led to the conclusion that the *trans* modes are feasible but less efficient than the *cis* configuration for recruiting Syk. These model predictions are supported by experimental evidence in mast cells transfected with chimeric receptors composed of WT and mutant versions of the γ cytoplasmic tail, fused to the transmembrane and extracellular domains of the Tac antigen (IL2α subunit). Crosslinking of the WT chimeric receptor with anti-Tac antibodies and secondary antibodies led to a robust calcium response, consistent with the ability of this construct to present phosphorylated ITAMs in *cis* to Syk. Chimeric receptors with alanine substitutions at Y64 or Y75, which can only recruit Syk via *trans* binding modes, were capable of eliciting measurable calcium responses. However, these responses were generally weak with a high percentage of non-responding cells. Our results are consistent with previous studies where γ-ITAM tyrosine mutants were expressed in γ-knockout bone marrow-derived mast cells (Yamashita et al., 2008). Since the β-ITAM is not engaged in the TT system, our results support the model that Syk recruitment to the γ-ITAM is the more prevalent interaction in the intact receptor. Calcium signaling was observed after TT-γ crosslinking in at least 20% of cells co-expressing of chimeric receptors with either Y64A or Y75A substitutions. These data suggest that the “diagonal” *trans* binding modes are more favorable for Syk recruitment than the other *trans* modes, a result predicted by the simulations reported in **Fig. 5**.

The argument that the *cis* configuration for Syk docking is by far the most efficient for signal transduction is also supported by model predictions that the stability of tandem SH2 domains docked onto paired pTyrs in the *cis* orientation is enhanced by novel contacts between “bystander” SH2 domains. These intriguing observations set the stage for experimental validation by site-directed mutagenesis, as well as comparisons with computational models of other tandem SH2 domain-containing proteins such as Zap-70, PTPN6, PTPN11 and PLCγ1 (Pluskey et al., 1995; Pei et al., 1996; Ji et al., 1999).

## MATERIALS AND METHODS

### Analytical model of multivalent interactions in WT Syk:FcεRIγ complex

The crystal structure of the Syk tandem SH2 domains bound to the ITAMs from a single CD3ε tail (PDB 1A81) (Fütterer et al., 1998) was used here to quantify the possible changes in Syk upon its binding to an immunoreceptor. In particular, the six complexes in the asymmetric unit cell of this crystal structure were compared after structural alignment of their N-SH2 in PyMOL (Schrödinger LLC, NY), which showed that the C-SH2 position varied within a range of 2.0 nm (*i.e.*, translational variability) and the C-SH2 rotation varied within a range of 18° (*i.e.*, orientational variability). The translational and orientational variabilities were used for computing the *V_acc_* (volume of accessible spherical shell) and *f_or_* (orientation factor) terms in equation (2) that gives the effective concentration *C_eff_* of unbound ITAM that unbound SH2 experiences upon binding of the other SH2:ITAM pair. This effective concentration can then be used in equation (1) to estimate the effective equilibrium association constant *K_overall_* for multivalent Syk:FcεRIγ, whose reciprocal is the effective equilibrium dissociation constant 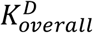 that can be compared with experimental measurements.

### System setup and unbiased MD simulations of Syk tandem SH2 and interdomain A

Chain A from PDB 1A81 was used as the initial conformation for performing unbiased MD simulations of Syk tandem SH2 and interdomain A. This chain contains the WT human sequence, and *in silico* mutagenesis was done using PyMOL to generate the WT murine sequence comprising residues 8-261. The initial conformation for the Syk Y130E mutant used this WT construct with the single phosphomimetic mutation done *in silico* using PyMOL. Acetyl and N-methyl neutral caps were added to the N- and C-termini of both constructs using PyMOL. All-atom topologies and parameters for both constructs were generated using the CHARMM36 force field (Klauda et al., 2010; Best et al., 2012). Solvent molecules were added using the TIP3P water model (Jorgensen et al., 1983) to fill a rhombic dodecahedral box around each construct, with a minimum distance of 1.2 nm from any protein atom to any edge of the simulation unit cell. Monovalent K^+^ and Cl^−^ ions were added to neutralize the system charge and to reach a physiological ionic strength of 150 mM.

The AMBER MD engine (version 16) that has been GPU-optimized for simulating explicit solvent systems (Salomon-Ferrer et al., 2012; Case et al., 2017) was used for running the unbiased MD simulations of the Syk WT and Y130E systems. Particle mesh Ewald (μME) electrostatics (Darden et al., 1993) were used along with Coulomb and Lennard Jones cut-offs of 1.2 nm and potential switching at 1.0 nm. Constant temperature was maintained at 310 K via Langevin dynamics (Pastor et al., 1988) with a collision frequency of 1.0 ps^−1^. Semi-isotropic pressure coupling was set for each system at 1 bar using a Monte Carlo barostat (Åqvist et al., 2004) with a relaxation time of 4.0 ps. Bonds containing hydrogen atoms were constrained using the SHAKE algorithm (Ryckaert et al., 1977). A hydrogen mass repartitioning approach (Feenstra et al., 1999) allowed the use of a 4-fs time step. Both systems were energy minimized using up to 10000 steps of steepest descent. Seven replicates of each system were then equilibrated, with each replicate having different initial velocities. Position restraints were applied to all protein heavy atoms during equilibration, which comprised 100 ps in the NVT ensemble and 10 ns in the NPT ensemble. Production runs (without position restraints) of 2 μs in the NPT ensemble were then performed for each replicate with configurations and energies saved every 200 ps. Two out of seven replicates were randomly selected for each system, and the simulations extended to 6 μs. Analyses were performed on the last 2/3 of each simulation. The distance reaction coordinate was measured between the Cα atoms of residue R21 and R174 that are part of the corresponding pTyr-binding sites on N-SH2 and C-SH2, respectively. The dihedral reaction coordinate was measured using the Cα atoms of R21 and L28 from N-SH2 with R174 and V181 from C-SH2, which belong to stable α-helices near the pTyr-binding site of each SH2 domain.

### System setup and REMD simulations of Syk interdomain A

The initial conformation for performing REMD simulations of Syk interdomain A was taken from the WT murine structure and comprised residues 115-159. As for the larger Syk construct (residues 8-261), the single phosphomimetic mutation Y130E was introduced *in silico* using PyMOL along with acetyl and N-methyl neutral caps on the N- and C-termini. Generation of all-atom topologies/parameters and the addition of solvent molecules and monovalent ions were performed as described earlier. The GROMACS MD engine (version 5.1.2) (Abraham et al., 2015) was used for running all simulations described in this section. Simulation parameters were similar as described earlier with a few exceptions. Pressure coupling was done using a Parrinello-Rahman barostat (Parrinello and Rahman, 1981), and bonds containing hydrogen atoms were constrained using the LINCS algorithm for the protein (Hess, 2008) and the SETTLE algorithm for waters (Miyamoto and Kollman, 1992). A 2-fs time step was used as hydrogen mass repartitioning was not performed for these systems.

Pilot unbiased MD runs were first run for the WT and Y130E mutant systems in order to determine the optimal temperature distribution. After energy minimization, a copy of each system was equilibrated at nine different temperatures equally spaced within the range 275-475 K. Energy minimization and equilibration were performed as described earlier for the unbiased simulations. Production runs (without position restraints) of 100 ns in the NVT ensemble were then performed with configurations and energies saved every 10 ps. The average energy and its standard deviation was then computed for each of the ten simulations for a particular system, and used as input for a REMD temperature scheduler (Garcia et al., 2006). This scheduler provided a temperature distribution comprising 63 temperatures within the range 275-475 K that would ensure a 20% exchange rate between all adjacent replicas in the REMD simulations. The same temperature distributions were given by this scheduler for the WT and Y130E mutant interdomain A systems.

These 63 temperatures were then used for running the REMD simulations. From the energy-minimized structures, a copy of each system was equilibrated at one of the 63 temperatures. All equilibration runs were performed as described earlier for the unbiased simulations. Production REMD runs (without position restraints) of 1.2 μs in the NVT ensemble were then performed for each replica with configurations and energies saved every 10 ps. Alternating exchanges between adjacent replicas were attempted every 2 ps, with swap acceptance or rejection determined by the Metropolis criterion. All analyses were done using the fifteen replicas with temperatures less than 310 K (excluding the 275 K replica that acted as a sink replica). Helicity for each of the fifteen replicas was monitored over the course of the entire trajectory, while subsequent processing and analyses used the last 500 ns of these trajectories.

### Hybrid MD/polymer model of multivalent interactions in WT and Y130E Syk:FcεRIγ complexes

The conformational ensembles for WT and Y130E mutant Syk interdomain A were each taken from the fifteen replicas described earlier for the REMD simulations. To generate the corresponding ensembles of WT and Y130E mutant Syk tandem SH2 and interdomain A, the N-SH2 (residues 8-116) and C-SH2 (residues 158-261) from the WT murine model of Syk were reattached *in silico* to every snapshot in the interdomain A ensembles using PyMOL. In particular, the atoms in the peptide bond between residues 115-116 were structurally aligned between N-SH2 and each interdomain A snapshot, and similarly for the atoms in the peptide bond between residues 158-159 to attach C-SH2. After filtering out of snapshots containing steric clashes between N-SH2 with interdomain A, C-SH2 with interdomain A, and N-SH2 with C-SH2, around 5% of the WT snapshots remained as physically viable models of Syk tandem SH2 and interdomain A, and similarly 5% of the Y130E mutant snapshots remained. The distance distribution between both SH2 domains for each system was obtained using the distance reaction coordinate described earlier.

These inter-SH2 distance distributions were used as the MD component in the hybrid MD/polymer model for estimating the effective concentration *C_eff_* in equation (3). The inter-ITAM-pTyr distance distribution comprised the polymer model component of this hybrid approach, and was generated using equation (4) that assumes each Syk molecule binds to single immunoreceptor tail in a *cis* mode. This equation requires a value for the peptide contour length, which represents the maximum/extended end-to-end length of the peptide as a function of the number of residues (*N*) (Zhou, 2001):

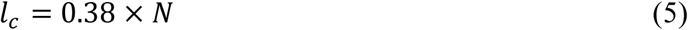

and a value for the peptide persistence length that gives the end-to-end length of the peptide beyond which a description of its behavior transitions from a flexible rod to a random walk (Rawat and Biswas, 2009):

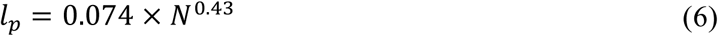

A value of 12 was used for *N* in equations (5) and (6), as this gives the length between the two pTyr residues in the FcεRIγ sequence (counting inclusive of both pTyr). *C_eff_* is then calculated in equation (3) by integrating over the products of the probabilities in the overlapping regions between both distributions for all possible distances (Van Valen et al., 2009; Sethi et al., 2011). *C_eff_* can then be used for getting the equilibrium dissociation constant 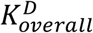 via equation (1) and getting the reciprocal.

### System setup and unbiased MD simulations of WT Syk:FcεRIγcomplexes with 2:2 stoichiometry

Although there are 24 possible permutations for combining four SH2 domains from two Syk molecules with four pTyrs from an FcεRIγ dimer, we found that only one *cis* and five *trans*modes can be built as physically viable structural models. The structure for the *cis* mode complex was obtained by first taking one of the Syk:CD3ε structures (chains A and B) from PDB 1A81 and performing *in silico* mutagenesis with PyMOL to get the WT murine Syk and FcεRIγ sequences, then placing two of these structures adjacent to each other. Each of the five *trans* structures were then obtained from the *cis* structure via a combination of manual translation and rotation operations in PyMOL. Backbone breaks/reseals and rotations were performed on the N-Cα and Cα-C bonds of only the FcεRIγ chain to keep the WT structure of Syk intact and to prevent accidental formation of peptide bonds with incorrect dihedral angles (*i.e.*, not 180°). Acetyl and N-methyl neutral caps were added to the N- and C-termini of all chains using PyMOL. Generation of all-atom topologies/parameters, addition of solvent molecules and monovalent ions, and unbiased MD simulations using GPU-optimized AMBER were performed as described earlier for the unbiased simulations of Syk tandem SH2 and interdomain A.

### Reagents and antibodies

Minimal essential medium (MEM) was purchased from Life Technologies (Grand Island, NY). Anti-Tac IgG was purchased from BioLegend (San Diego, CA) and the anti-mouse IgG purchased from Jackson ImmunoResearch Laboratories, Inc. (West Grove, PA). Transfection materials solution L and Amaxa^TM^ Nucleofector^TM^ were purchased from Lonza (Basel, Switzerland). Fura-2AM was from Molecular Probes (Eugene, OR).

### Cell culture and transfection

RBL-2H3 cells (Metzger et al., 1986; Wilson et al., 2000) were cultured in MEM supplemented with 10% heat-inactivated fetal bovine serum, puromycin, and l-glutamine. Cell health and behavior was confirmed by checking IgE binding and degranulation response after each thaw and cells were used for no more than 10 passages. Transfections were conducted using the Amaxa system (Lonza) with Solution L and Program T-20. Transfected cells were then plated in 8-well tissue culture plates in MEM supplemented with 15% heat inactivated fetal bovine serum, 50 U/ml penicillin, 0.05 mg/ml streptomycin, and 2 mM L-glutamine. The expression vector for the chimeric TT-γ receptors has been previously described (Letourneur and Klausner, 1991). For all microscopy experiments, cells were plated overnight in 8-well Lab-Tek (Nunc) chambers (ThermoFisher Scintific) at a density of 50,000 cell/well.

### TT-γ crosslinking and calcium measurements

Calcium measurements were performed as previously described (Schwartz et al., 2015). Fura-2AM loading (2 nM in Hanks buffer) was performed for 30 minutes at room temperature with 1 μg/ml anti-Tac IgG added for the final 10 min. After Fura-2AM loading, cells were washed and transferred to the Olympus IX71 where cells were maintained at 35 °C using an objective heater (Bioptechs). Cells were activated by crosslinking with anti-mouse IgG (25 μg/mL), added at 30 seconds during a total of 5 minutes of imaging. The ratio of fluorescence emission with excitation at 350/380 nm was calculated for each cell (5-10 per field of view) over time to assess calcium release.

## ACKNOWLEDGMENTS

This work was supported by National Institutes of Health (NIH) grants to the New Mexico Spatiotemporal Modeling Center (P50GM085273 for B.S.W.; Department of Energy through contract DE-AC5206NA25396 for S.G. and T.T.; R01GM100114 for D.S.L.). T.T. is also supported by the Center for Nonlinear Studies (CNLS) at the Los Alamos National Laboratory (LANL). Computing resources were made available through Extreme Science and Engineering Discovery Environment (XSEDE) Allocation MCB170148, which is supported by National Science Foundation (NSF) grant number ACI-1548562, and through LANL Institutional Computing. We thank Shayna Lucero for assistance with cell culture and Dr. Cedric Cleyrat for generation of the TT-γ mutants, and gratefully acknowledge use of the University of New Mexico Cancer Center fluorescence microscopy facility, as well as NIH-NCI support via P30CA118100 for this core.

